# ANALYSIS OF THE PHOSPHORYLATION NETWORKS CHARACTERIZING DISTINCT PHENOTYPIC STATES IN GLIOBLASTOMA CELL POPULATIONS

**DOI:** 10.1101/2025.08.22.671611

**Authors:** Januka Khanal, Chunxiao Ren, Wenxue Li, Tatsuya Matsubara, Jihoon Lee, Deok-Ho Kim, Jennifer Moliterno, Murat Gunel, Sergei Kotelnikov, Stan Xiaogang Li, Ernest Glukhov, Dima Kozakov, Yansheng Liu, Andre Levchenko

**Affiliations:** Systems Biology Institute, Yale University, West Haven, CT 06516; Biomedical Engineering Department, Yale School of Engineering, New Haven, CT 06511; Molecular Biophysics and Biochemistry, Yale University, New Haven, CT 06511; Cancer Biology Institute, Yale University, West Haven, CT 06511; Department of Pharmacology, Yale School of Medicine, New Haven, CT 06511; Department of Biomedical Engineering, Johns Hopkins University School of Medicine, Baltimore, MD 21205; Department of Neurosurgery, Yale School of Medicine, New Haven, CT 06511; Department of Applied Mathematics and Statistics, Stony Brook University, Stony Brook, NY, USA; Laufer Center for Physical and Quantitative Biology, Stony Brook University, Stony Brook, NY, USA; Oden Institute for Computational Engineering and Sciences, The University of Texas at Austin, Austin, TX, USA; Department of Molecular Biosciences, The University of Texas at Austin, Austin, TX, USA

## Abstract

Cancer cells, including the most aggressive brain cancer, glioblastoma, can exhibit a high degree of phenotypic plasticity. However, it is not well understood whether and how distinct phenotypic states might be associated with or driven by specific signaling processes. Here, we use a novel approach to the identification of phenotype-specific signaling networks in cells undergoing the Go-or-Grow switch in populations of GBM cell lines. We find that the transition to invasive spread may be associated with the onset of DNA damage response, cell cycle arrest, and activation of the AKT-mTOR signaling. We reconstructed the large-scale signaling networks, revealing integration of diverse pathways and possible feedback interactions mediating phenotypic stability. We further show that phosphorylation outcomes may be preferentially mediated by 14-3-3 proteins, previously implicated in stress response and control of the cell cycle. Finally, we demonstrate that the phenotype-specific phospho-site signatures have predictive power for disease-free patient survival. This study paves the way for a more comprehensive understanding of phenotypic plasticity in GBM and other aggressive cancers.

## INTRODUCTION

Glioblastoma (GBM), classified as the Grade IV glioma by WHO, is the most common and deadliest form of brain cancer. This cancer has an extremely poor prognosis, with a Median survival of approximately 15 months post-diagnosis and a progression-free survival of 5–6 months following application of the standard of care^1,2^. The high mortality in GBM is associated with limited treatment options, which themselves may be due to high genetic and phenotypic heterogeneity found within the tumors ^3–5^. Historically, heterogeneous mutational loads across the tumor cell populations have been attributed to genetic mutations and dynamic clonal selection. However, it has become clear that even the cells with identical genetic makeup can display a high degree of phenotypic heterogeneity and plasticity.^6–8^ Examples of such phenotypic plasticity include switches between quiescent, proliferative, and highly invasive and migratory cell states (the so-called ‘go-or-grow’ phenotypic switch) or switches to drug- or radiation-resistant states during therapeutic interventions. Such plastic responses can enable subpopulations of cancer cells to evade surgical resection and targeted treatment, leading to relapse and further disease progression.

Importantly, phenotype switching has also been observed for cells and tumors that have been characterized into different subtypes based on their genetic, transcriptomic, and metabolic states. In particular, single-cell transcriptomic analysis not only suggested the existence of four cellular states (neural progenitor–like (NPC-like), oligodendrocyte progenitor–like (OPC-like), astrocyte-like (AC-like), and mesenchymal-like (MES-like), but also suggested that they are not stable and cells can switch between these states over time.^8^ Recent proteogenomic studies propose multi-omics classifications, incorporating protein and phosphoprotein profiles have confirmed that phenotypic plasticity can be observed for cells characterized by distinct states of their regulatory network^3,7^. In combination, these studies suggest that phenotypic plasticity characterized by either the multi-omics assessment or direct functional analysis of cellular phenotypes can be a substantial challenge to the treatment of GBM and potentially other highly aggressive cancers.

The mechanisms underlying phenotypic plasticity are still poorly understood, although they likely depend on the increased epigenetic dysregulation observed in cancer cells.^9,10^ Furthermore, the dynamics of the transitions between different states are not characterized, and it is not clear whether it may depend on the underlying genetic and epigenetic background of the cancer cells. As changes in the cell micro-environment may trigger the phenotypic transitions, the more plastic cells in different initial states might quickly conform to the stimulus by fast adaptive switching, leading to a progressively lower phenotypic heterogeneity. On the other hand, in populations of less plastic cells, the heterogeneity may persist and even potentially be amplified by differential cell responses to the stimulus. Characterization of such differences in the degree of cell plasticity may help develop more personalized approaches to cancer treatment.

In this and companion studies, we address the questions of what may specify different phenotypic states and whether they may be affected by the different genetic background of GBM cells. We perform this analysis for the clinically relevant phenotypic switch associated with the triggering of the invasive spread and dissemination of cancer cells from the primary tumor site. These invasive cells can be difficult to remove during surgery. They can display particularly high levels of resistance to chemo- and radiation therapies ^11,12,13,14^, seeding the growth of the recurrent tumors. We use a previously characterized experimental platform mimicking invasive cells spread (which is predictive of clinical outcomes when used with primary patient-derived cells^15^ coupled with a multi-omics analysis to identify and characterize phenotypically distinct, migratory, and proliferative GBM cell sub-populations. Here, we focus on the phospho-proteomic analysis of more migratory and more proliferative phenotypic states isolated by the phenotypic filtering enabled by this platform. We extensively characterized the long-term dynamics of cell response to the extracellular matrix (ECM) cue triggering invasive cell spread within this experimental analysis, finding that the cell migratory state is enabled by the activation of a set of signaling networks, distinct and likely inhibitory of the pro-proliferative signaling. These networks mediate the response to cellular stresses, trigger G2M checkpoint activation, increase mTOR signaling, and glycolytic metabolism. We further demonstrate that the phosphoproteomic signature can be highly predictive of disease-free survival of GBM patients.

## RESULTS

### Phenotype Filtering using the Rapid Assessment of Cell phenotypic Extremes (RACE) assay can sort GBM cells into migratory and non-migratory subpopulations, and examine the degree of phenotypic plasticity

Previously, we showed that an ECM-mimicking cell adhesion substrate promotes the transition of primary and serum-cultured GBM cells into a pro-migratory state highly resembling migratory behavior in the brain tissue, in contrast to the usual conditions of culturing cells on flat and hard plastic and glass surfaces ^15^. Furthermore, we showed for a large set of cancerous and normal cells of different types that this ECM-mimetic micro-environment can promote epithelial-to-mesenchymal transition and more aggressive cell spread (REF). Here, we used this experimental platform as both a pro-migratory cue and a natural tool to separate cellular phenotypes. Indeed, a highly directed movement of the subset of more migratory cells can allow them to move away from the rest of the cell population, similar to the invasive cell spread in vivo. We can then capture and isolate these highly migratory cells, as well as the cells showing the lowest motility and characterize them as cells adopting distinct degrees of the migratory phenotype. Furthermore, this process can be repeated over multiple rounds, with both the ‘fast’ and ‘slow’ cell sub-populations again sorting themselves and potentially enriching the phenotypic state. The enrichment is expected to work particularly well if the cells have a relatively low degree of phenotypic plasticity and can thus retain their phenotypic state with high temporal persistence.

On the other hand, relatively more plastic cells would be expected to not retain the specific phenotypic state over a long period of time and might be more affected by the pro-migratory cue, potentially forcing all cells to adopt a cue-dependent phenotypic state. This analysis, termed Cell phenotypic Extremes (RACE) assay, thus allows us to have a working definition of the degree of plasticity and use this analysis on cells with different genetic backgrounds. Here, we focused on the characterization of the plasticity and (phosphor-)proteomic analysis of phenotype specific networks, following the pipeline shown in Fig. 1A.

**Figure 1:**
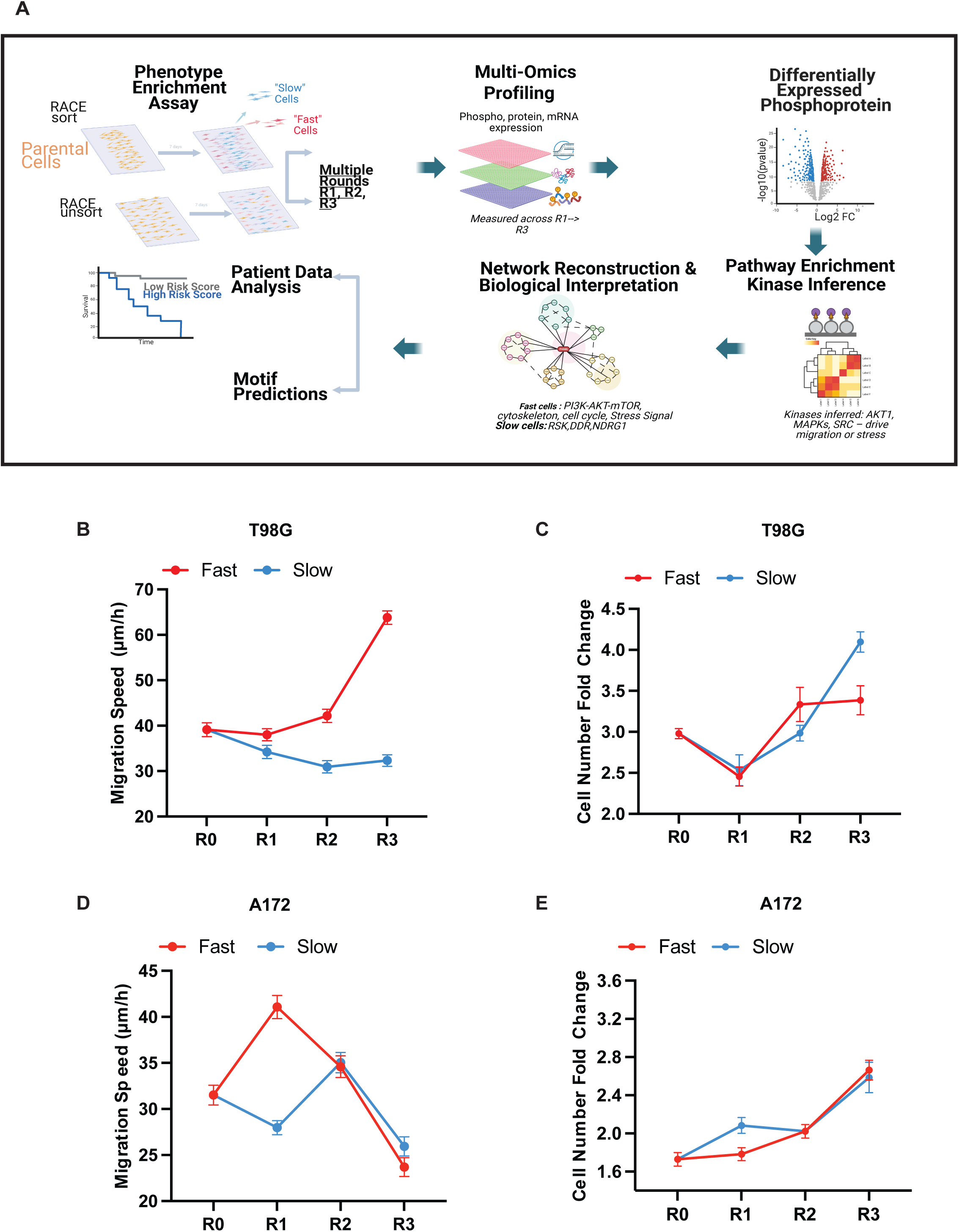
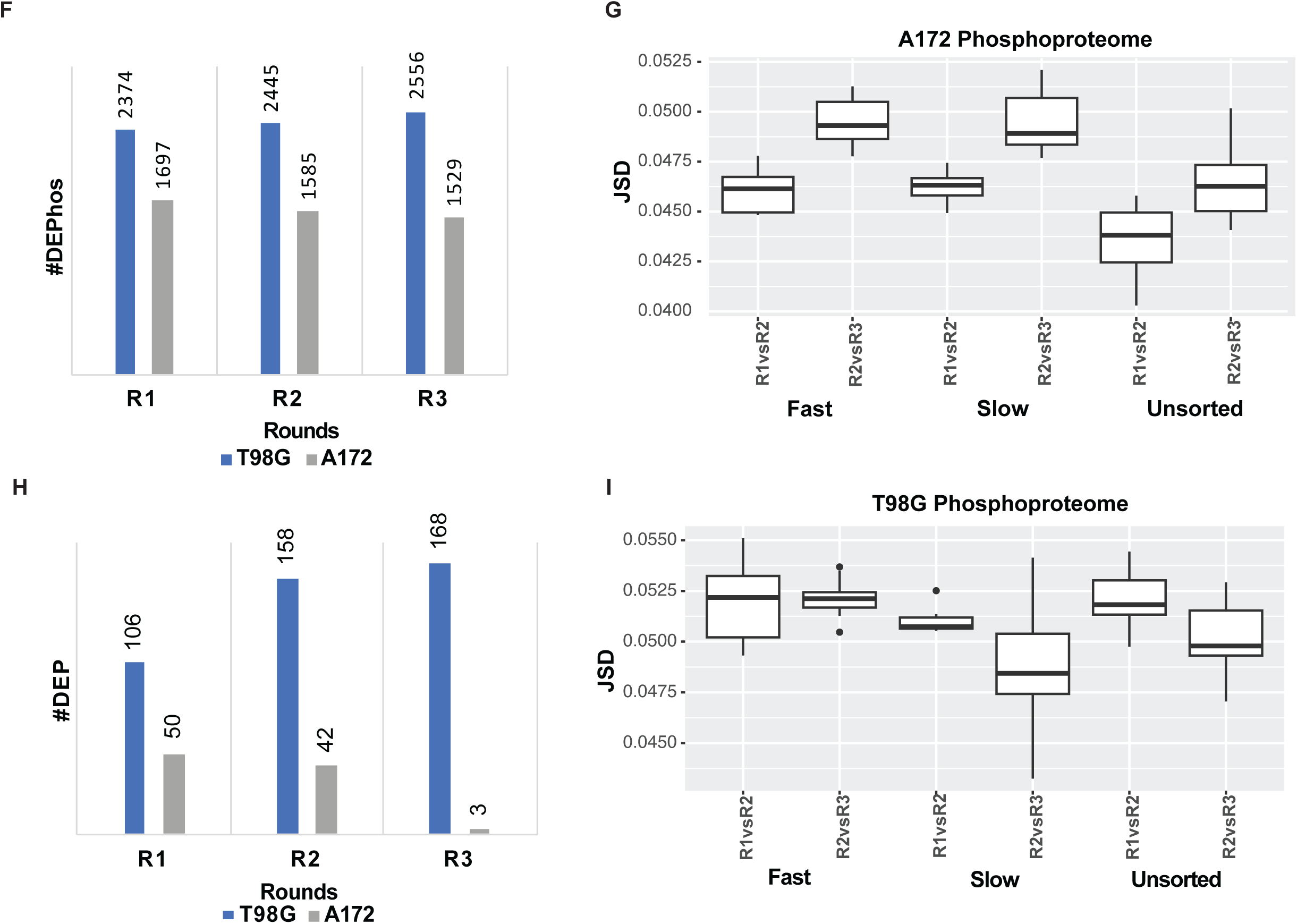
Phenotypic characterization of GBM subpopulation(A) Pipeline of the research (B and D) Measurement of cell migration speed in μm/hr across rounds of RACE (C and E). Measurement of cell proliferation rate across rounds of RACE. (F and H) Number of Differentially Expressed Phosphosites (DEPhos) and Proteins (DEP) (log2FC>|1| and p value < 0.05) between fast and slow cells over rounds. Numbers on top of the bar graph represent the exact number of DEPhos and DEP for that particular round. (G and I) Jensen Shannon Divergence score for T98G and A172 subpopulations. P0: Passage 0: Sorted cells collected right after the RACE assay. P1: Sorted cells collected after passing once. R0: Cells that were plated on 2D glass plates.

We focused on analyzing two GBM cell lines that have distinct commonly observed mutational profiles, with key results that were further validated vs. a large human patient cohort. These cell lines (A172 and T98G) have been previously shown to retain the key characteristics of clinical GBM ^16^ Among the key distinguishing genomic features, A172 cells are wild type for TP53, whereas T98G cells harbor a missense mutation in this gene. In contrast, A172 cells are null for expression of PTEN, and T98G cells express a wild-type copy of this gene. Both cell lines displayed migratory and proliferative responses on the RACE platform. Interestingly, as further detailed in the companion study, we found highly divergent trends in the dynamics of phenotypic responses displayed by cell sub-populations over multiple over multiple rounds of the RACE assay. In T98G cells, the filtered sub-population of Fast cells showed a progressive enrichment of highly migratory cells and a decrease in the average cell proliferation. In contrast, the Slow cells showed the opposite trend. Overall, there was a progressive divergence of phenotypic cell characteristics between the Fast and Slow sub-populations. In a stark contrast to this behavior, in A172 cells, the initially divergent phenotypic features observed most clearly in Fast vs.Slow cells after the first round, gradually converged with both cell sub-populations, revealing similar migratory and proliferative behavior by the third RACE round. In agreement with our working definition of cell plasticity, these divergent characteristics suggested that A172 cells were more plastic than T98G cells, and that transitioned to the more uniform phenotypic state driven by the biomechanical cue embedded in the RACE assay, whereas T98G cells retained the memory of their initial phenotypic states. The results also suggested that Slow cells in both cell lines are more proliferative, supporting the Go-or-Grow dichotomy.

To examine if divergent phenotypic dynamics are also supported on the level of phosphorylation-based post-translational modification networks, we performed the proteomic and phospho-proteomic analysis of Fast and Slow cell sub-populations for both cell lines over three rounds of RACE. The samples were measured by the data-independent acquisition (DIA) MS method as described previously ^3-5^. In a striking parallel to the phenotypic analysis, we found that there was a progressive increase for T98G cells and a progressive decrease for A172 cells, in the number of both differentially expressed proteins (DEPs) and differentially expressed phospho-sites in the Fast and Slow cell sub-population over three RACE rounds. This result was consistent with a similar dynamics of divergence in the number of differentially expressed genes for sub-populations of T98G cells and convergence of this number in A172 cells, as reported in the companion study. These results supported a gradual enrichment of cells displaying distinct phenotypes by the RACE assay in T98G cells, and a gradual convergence of both phenotypic and regulatory network states in the A172 cells.

### Molecular and pathway-level characterization of the phenotype-specific phospho-proteomic profiles associated with the migratory and proliferative phenotypes

We next examined the phospho-proteomic profiles characterizing the Fast and Slow phenotypic states. We found that 103 unique phospho-sites for T98G (Fig. 2A) and 37 for A172 (Fig. 2E) remained differentially expressed between fast and slow cells across all rounds of the RACE assay. We next sought to explore further the changes in the expression of these phospho-sites across rounds and to annotate their potential functional roles. Specifically, we used supervised K-means clustering to identify groups of phospho-sites whose expression might be correlated over the rounds. This analysis revealed four general dynamical trends one expects to see in the data: phospho-site groups with steady levels that are either higher across rounds for Fast or for Slow cell sub-populations, and phospho-site groups showing a relative increase or relative decrease in the relative phospho-site abundance in Fast vs Slow cells. We indeed observed that K-means clustering identified these four dynamic trends as different clusters for T98G cells(Fig. 2B and 2C). For A172 cells, only two clusters were distinguishable with more mixed internal dynamics, which, nevertheless, on average reflected the gradually convergent network activity (Fig. 2E and 2G).

**Figure 2:**
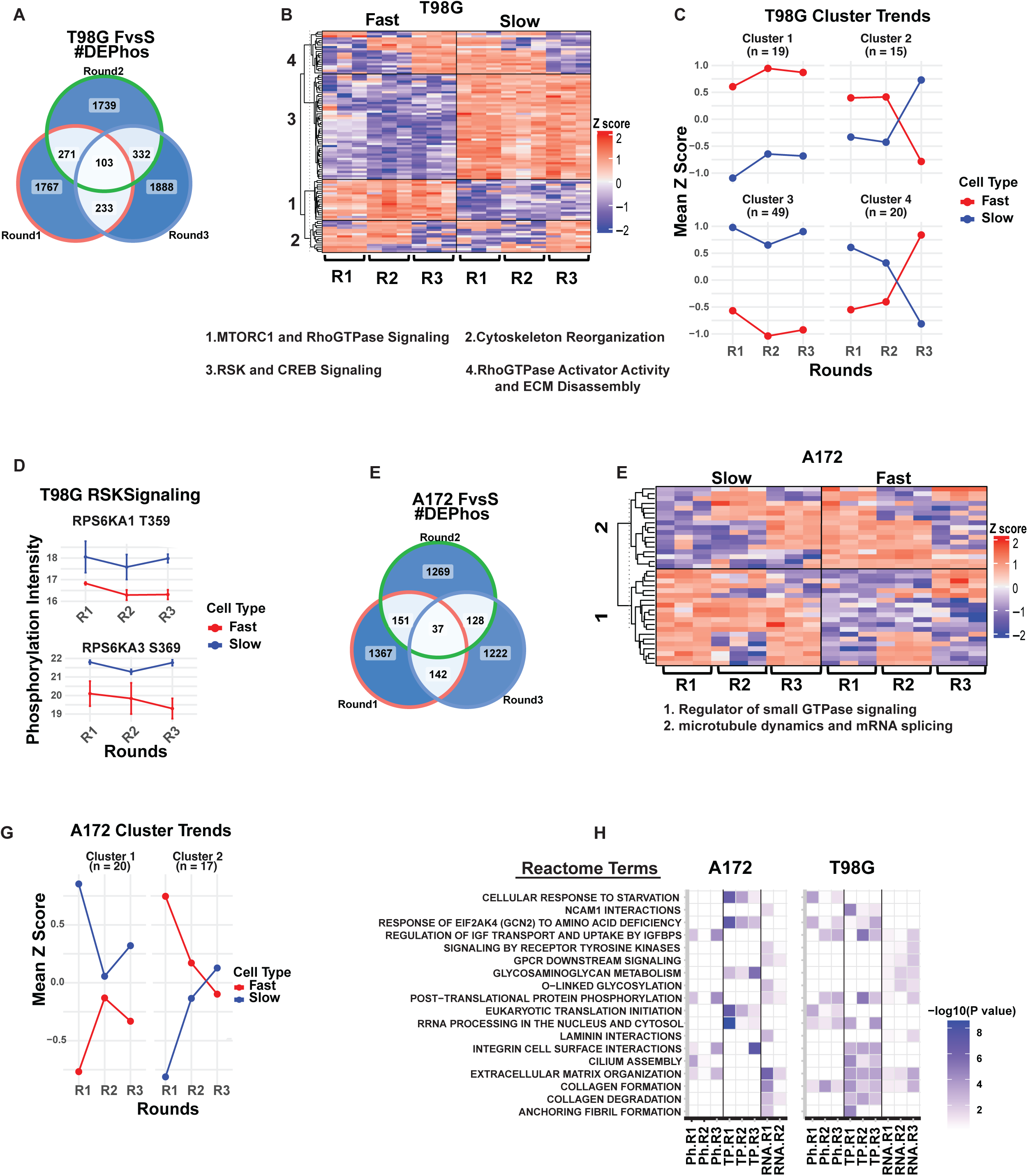
Dynamic and Pathways enrichment of sustained Differentially Expressed Phosphophosites (DEPhos). (A and E)Venn diagram for the number of DEPhos between fast and slow (FvsS) through Rounds. (B and F) Heat map of K-means clusters of sustained DEPhos for T98G and A172, respectively. (C and G) Average mean Z-score dynamics of the DEPhos in B and F. RSKs Phosphorylation dynamics trends across rounds. (H) Reactome enrichment overlaps between two cell lines across Phosphoproteomics (Ph), Proteomics (TP), and Transcriptomics(RNA).

To gain a better insight into the functional significance of these results, we mapped the phospho-sites to the specific proteins to assess their putative involvement in the Fast/Slow phenotypic dichotomy. First, we analyzed the four clusters of T98G cells. We found that (as revealed by cluster 1 phospho-sites) proteins responsible for RNA processing (Fig. S5), ECM disassembly (e.g., MAP2, LCP1), and cell adhesion/cytoskeletal signaling (e.g., FRMD5 and TRIP6) (Fig. S1A), were consistently highly up-regulated in Fast cells. On the other hand, (as revealed by cluster 3 phospho-sites), phosphorylation of other cytoskeletal proteins, such as MTUS1, KIF16B, ABLIM3, IQGAP2, FRMD4B, and LIMCH1, was up-regulated in the Slow cells (Fig. S1C). Furthermore, components of two key signaling pathways – ERK-RSK (e.g., RPS6KA1 and RPS6KA3) and PKA-CREB (e.g., PRKAR2B, AKAP6) also displayed consistently increased phosphorylation in Slow cells (Fig. 2D and S1F). Notably, the up-regulation of RSK phosphorylation in Slow cells on S359/S369 has been associated with the increase in its activity, which may be further enhanced by the relative increases in the mRNA and protein levels of the isoforms of this kinase observed in Slow vs. Fast cells here and the companion study.

The analysis of clusters 4 and 2 revealed the gradual enrichment of phosphorylated proteins in the Fast and Slow cells, respectively, over the course of multiple RACE rounds. In particular, we observed an increase in the relative phosphorylation of stress response proteins (MLST8, HSPA8 (Fig.S1E)), components of beta-catenin signaling (e.g., BCL9L, KMT2D), and mTOR signaling (e.g., LARP1, MLST8 (Fig.S1E)) in Fast T98G cells. The proteins whose phosphorylation showed a relative increase in Slow cells included PHLDB1, implicated in GBM proliferation and other gene products with possible roles in GBM progression (e.g., CORO1A and PDLIM5)^17^(Fig. S1B). Of particular interest was enhanced phosphorylation of ZMYM3, a stress response protein controlling chromatin modification and DNA repair (Fig. S2B).^18^ Epigenetic regulation by this and other factors may underlie cellular plasticity by controlling gene expression programs relevant to distinct phenotypes^10^.

Analysis of clustering results performed for A172 cells, revealed the greatest difference in the levels of protein phosphorylation between Fast and Slow cells in Round 1 of RACE, with higher phosphorylation in Fast cells of the Rho GTPase and cytoskeletal signaling (e.g., ARHGEF2, DOCK4, ARHGAP1) (Fig. S1J) and for lipid metabolism/cholesterol regulation (e.g., FASN, CHD9, SREBP). The proteins showing relatively higher phosphorylation levels in the Slow cells included regulators of cytoskeletal and microtubule dynamics (e.g., MAP1A, TPX2, MAP7D3), RNA splicing (PRPF31, SNRNP200, TRA2A)(Fig.S1G), and cell proliferation (e.g., CENPT) (Fig. S1I). We again detected increased relative phosphorylation of ZMYM3, suggesting that regulation of this protein may be a conserved feature in the phenotypic control of GBM cells.

We next took advantage of the availability of multi-omic (phospho-proteomic, proteomic, and transcriptomic) data for the matched samples collected for both cell lines across multiple RACE rounds. We analyzed these data in terms of the reactome enrichment terms to examine whether there was consistency between differential mRNA and protein expression and phosphorylation levels in Fast and Slow cells across the rounds of the RACE assay. We found that although there was an overlap between these data for both cell lines, particularly in regulation of integrin-mediated signaling and ECM regulation, likely due to the effect of the bio-mechanical input associated with the RACE platform, there was also a substantial degree of divergence between the results, indicating that each modality of the multi-omic analysis can provide information that may not gleaned from the other modalities (e.g., the post-translational modifications revealed by the phosphoproteomics analysis may not be uniformly correlated with the gene and protein expression levels). Nevertheless, there was a consistent observation of an increase in the differentially expressed genes (DEGs) and the corresponding reactions revealed by the mRNA data analysis in Fast vs. Slow cells over multiple rounds for T98G cells and a decrease of DEGs for A172 cells, consistent with a similar dynamics of differentially expressed phospho-sites and phosphorylated proteins, reported above. This result suggested that there is a direct link between transcriptional control and the signaling networks regulating it.

To further address the putative role of phosphorylation in transcriptional regulation of the Fast and Slow cell sub-populations, we analyzed the transcription factors inferred as differentially activated based on the enrichment of their previously implicated target genes (see the companion study for details). Strikingly, we found that only a tiny fraction of these factors showed differential expression between Fast and Slow cells in both cell lines, both on the mRNA and protein levels (Fig. 4A and 4S1). This lack of detected expression difference was not due to the low levels of expression of these factors, as the TPM counts for their mRNA levels were comparable to those of other genes in the dataset (Fig.4D and 4E). However, we found that the inferred TFs were differentially phosphorylated in a phenotype-specific fashion in both cell lines(Fig. 4B and 4C). In agreement with prior results, the relative phosphorylation differences between Fast and Slow cells progressively increased over RACE rounds for T98G cells and decreased for A172 cells. These results suggested that protein phosphorylation, instead of changes in gene or protein expression, is a key regulator of differential gene programs activated in the GBM cell lines analyzed here.

**Figure 3:**
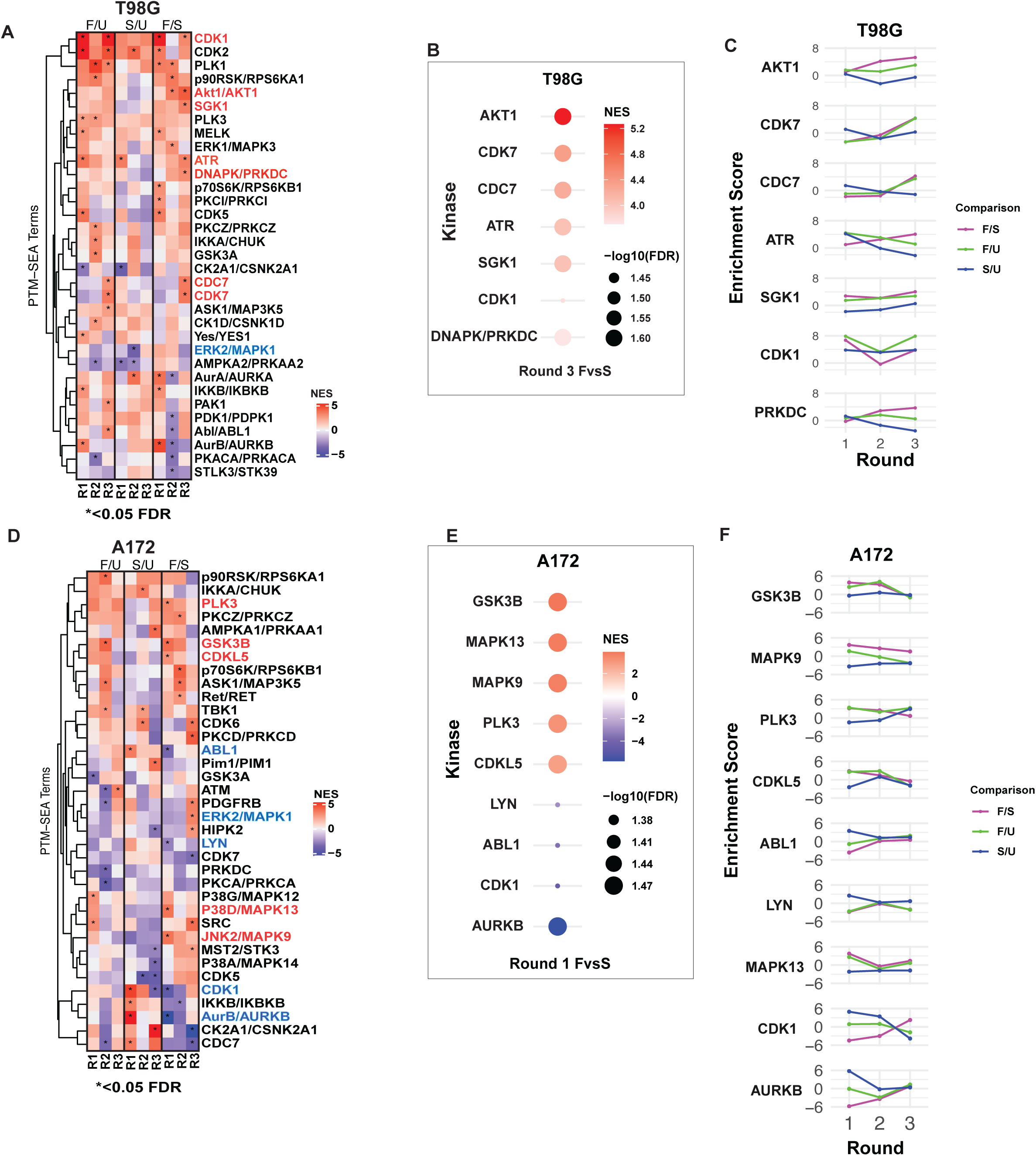
Inferred Kinases and the Dynamics: (A and D): Kinase activity inferred through ranked-based enrichment analysis, PTM-SEA. KINASE-PSP are the kinases that are inferred based on the phosphositeplus database. NES: Net enrichment score. *: significant enrichment between the groups compared (adj p-value<0.05). F/U column: Fast cells are compared to Unsorted cells. S/U column: Slow cells are compared to Unsorted Cells. F/S: Fast Cells are compared to slow cells. (B) T98G: Kinases with significant enrichment scores between fast and slow in Round 3. (E) A172: Kinases with significant enrichment scores between fast and slow in Round 1. (C and F) Dynamics of the kinases listed in B and C through the rounds of RACE.

**Figure 4:**
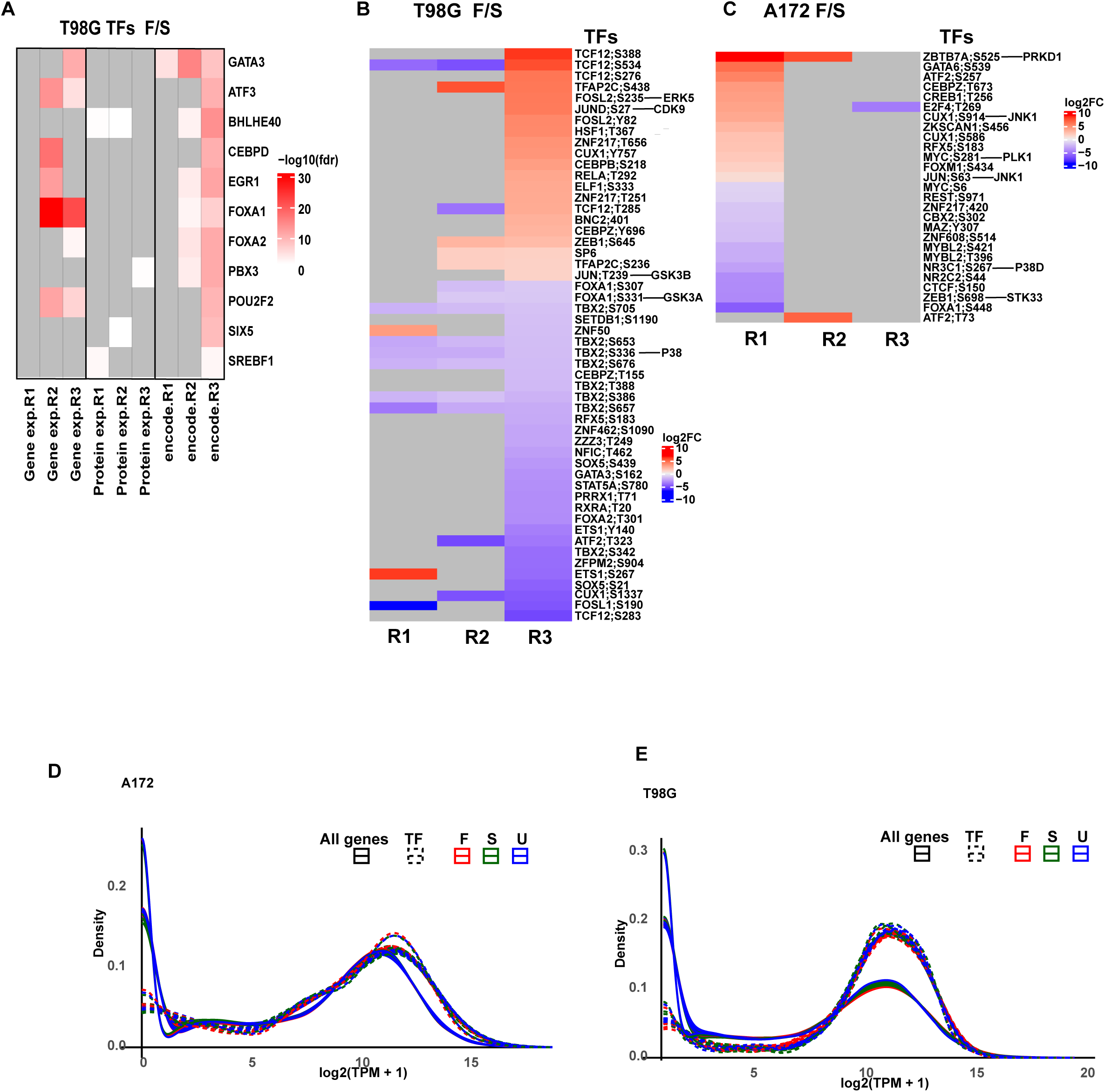
Phosphorylation of transcription factors dominates the differential gene expression regulation between fast and slow cells. (A) –log10(fdr) of the encode inferred (encode) Transcription factors(TFs), and their expression at the mRNA (Gene.exp) and protein (protein.exp) level. (B and C): Heat map of log2FC of Encode inferred Transcription factors that are significantly differentially phosphorylated through the rounds of RACE assay. The kinases regulating the particular TFs’ sites are shown on the right next to phosphosites. (D and E): The density plot of the transcript per million (TPM) counts for transcription factors (dashed line) against all genes in fast, slow, and unsorted populations (solid line).

### The kinase-substrate enrichment analysis suggests the key importance of several kinases in controlling the migratory and proliferative phenotypic states in GBM cells

Mammalian kinases have multiple substates and associated phospho-sites. Even though the precise enzyme-substrate matching is still lacking for the human kinome, and this task can be complicated by the ability of multiple kinases to phosphorylate the same phospho-sites, we attempted to reconstruct the subset of high-confidence inferences of the kinases involved in controlling phospho-sites enriched in Fast and Slow cells in GBM cell lines. In particular, we performed PTM-SEA (Post-Translational Modification Signature Enrichment Analysis)^19^ using Phosphosite Plus Data Base(PSP-DB) ^20^. The results were expressed as the normalized enrichment score (NES), and we focused on the significantly enriched kinases using a p-value cutoff of < 0.05. The NES values were examined for both cell lines across multiple RACE rounds.

Examining the data for T98G cells, we found that the following kinases passed the significance criteria for enrichment in Fast vs Slow cells: SGK1, AKT1, ATR, DNA-PK, CDC7, CDK7, CDK1 (Fig. 3A and 3B) in Round 3, at which Fast and Slow cells show the largest phenotypic differences. All these kinases, except CDK1, displayed progressive enrichment in Fast cells compared to Slow cells over the course of the RACE rounds (Fig. 3C). We noted that these results were consistent with our prior observations of mTOR and stress response signaling pathways in Fast vs. Slow cells. Indeed, both AKT1 and SGK1 can activate mTOR downstream of PI3K^21,22^, and both ATR and DNA-PK have been strongly implicated as key regulators of the DNA damage response^23^. Although CDK7 and CDC7 have the canonical roles as regulators of progression through the cell cycle, they can also control the DNA damage response and the corresponding cell cycle checkpoints, which may reconcile their increased activity with the relative decrease of proliferation in Fast cells^24,25^. Related to this, we found that the inferred CDK1 activity increases in both Fast and Slow cells relative to the control of the cells that were cultured on the RACE platform but not sorted into the Fast and Slow sub-populations (unsorted cells). CDK1 can be up-regulated both in the context of cell cycle progression and the spindle assembly checkpoint^26,27^. These results suggest that the interpretation of the cell cycle regulating kinase function can be complex, and their enrichment may be associated with both proliferative and cell cycle arrested cell states.

In A172 cells, the inferred kinase activity in Fast vs. Slow cells was up-regulated for JNK2 (MAPK9), p38D (MAPK13), PLK3, and CDKL5, whereas the activities of Abl, AURKB, and CDK1 were relatively increased in Slow cells(Fig.3D and Fig. 3E) in Round 1 at which Fast and Slow cells show the most distinct dichotomy. Over multiple RACE rounds, we found that kinases with positive NES decreases (enriched in Fast cells) displayed a reduction of the NES values by Round 3, and kinases with negative NES increased the NES value by Round 3(Fig. 3F). These results indicated up-regulation of stress response kinases that, though distinct from those up-regulated in the Fast T98G cells, can nevertheless represent an alternative mechanism of responding to the DNA damage stress. Indeed, p38 signaling is activated downstream of ATR and ATM in response to DNA damage stress^23^,^28^. The companion study provided further evidence for a relative increase in activation of p53-mediated response in Fast A172 cells, further suggesting that DNA damage response up-regulation occurs in these cells. The roles of PLK3 and CDKL5 can be related to the control of cell migration, particularly for the brain-derived cells, and may be specific to A172 cells. The up-regulation of inferred activity of CDK1, and particularly Aurora kinase B (AURKB) in Slow A172 cells was consistent with the higher proliferation rate of these sub-population. These results were supported by the analysis of the proteomic profiles in Slow vs Fast A172 Cells, suggesting strong up- regulation of the cell cycle regulator APC/C, based on enrichment of its enzymatic targets in Slow A172 cells reported in the companion study.

Overall, the results of our analysis suggest up-regulation of a specific subset of kinases associated with the Fast and Slow phenotypes. In particular, the results suggested active involvement of stress response and cell cycle-associated kinases in defining these phenotypic states and regulating the associated Go-or-Grow phenotypic switch.

### Network Reconstruction Using Integrated Multi-Omics Data Suggests Cell Line-Specific And Overlapping Phenotype Controlling Mechanisms in GBM Cells

A key challenge of the present-day systems-level reconstruction of phosphorylation-mediated signaling mechanisms is the fragmented nature of network analysis and the lack of annotation of the functional consequences of phosphorylation events. This work explores whether results from a high-throughput multi-omics analysis, such as those presented here, can be used to infer a self-consistent, non-contradictory network of both measured and inferred connections between signaling nodes. Specifically, the goal is to determine whether an unbiased reconstruction of phenotype-specific regulatory networks is feasible. To address this, a diverse set of computational tools and analytical methods was employed, as described below.

To explore the signaling mechanisms driving the migratory phenotypes, we constructed a signaling network model for each round of the RACE assay based on the differential activation of kinases and phosphoprotein expression. As previously noted, T98G at Round 3 (R3) and A172 at Round 1 (R1) exhibit the most pronounced differences in inferred kinase activity and cellular phenotypes such as speed and proliferation between the Fast and Slow states. It is therefore expected that these rounds will also display the most pronounced differences at the level of signaling network architecture. Thus, we investigated the signaling network at these rounds by reconstructing a signaling network using PHONEMeS (PHOsphorylation NEtworks for Mass Spectrometry)^29^, and enhancing the resulting unsigned network through the addition of regulatory edges between phosphoproteins, transcription factors, and protein and mRNA abundance. In particular, we implemented the upside-down model of PHONEMeS-ILP (PHONEMeS-UD), which enables bidirectional inference of kinase activity by considering both upstream and downstream signaling states. In PHONEMeS-UD, the consistently up- and down-regulated kinases inferred from phosphoproteomics data were integrated as putative perturbed nodes, allowing the model to infer both kinase activation cascades and phosphorylation-dependent regulatory mechanisms. To refine the inferred networks and enhance their interpretability, we further added the edges representing inhibition and stimulation when available using the PhosphoSitePlus database (PSP-DB). To prevent contradictions in inferred signaling pathways, we also introduced logical consistency constraints. These constraints ensured that the inferred signaling mechanisms did not contradict known biochemical regulation, such as the incorrect inference of activating phosphorylation leading to kinase inhibition. Further, we added missing edges between the nodes in the network and master regulators of the input kinases in the network based on prior knowledge and literature.

Finally, to connect phosphoproteomics anlysis with inference of gene regulatory mechanisms, we integrated transcription factors (TFs) inferred from RNA-seq data into the phospho-signaling network. This integration was performed using both a bottom-up approach (linking TFs to upstream kinases) and a top-down approach (tracing kinase-driven regulation of TFs), resulting in a multi-layered regulatory network. The kinases regulating the TFs (bottom-up approach) were identified from the PSP database and the Kinase Library. Additionally, we incorporated mRNA and total protein expression data to account for transcriptional and translational regulation of the differentially phosphorylated proteins. Specifically, we included genes with differential expression in total proteomics (adjusted p-value < 0.05 and |log₂FC| > 1) and mRNA expression (adjusted p-value < 0.05) to annotate the network further.

The resulting network reconstructions (Fig. 5B and 5C) put the prior phospho-site, kinase, and reactome results into the context of a larger interconnected signaling network. Unlike conventional network representation, the inputs in these network reconstructions were from the kinases whose activity showed clear differential activity in our prior analysis. These results indicated that the network reconstruction is indeed feasible and that one can generate a putative representation of both kinase crosstalk and combined regulation of signaling targets and the corresponding gene/protein expression regulation. Furthermore, this analysis permitted tentative separation of distinct signaling sub-networks or modules, which contained sub-networks that, though varying in details between the cell lines, suggested a similar overall mechanism of separation into the Go and Grow phenotypes in GBM cells. In particular, the analysis fleshed out the interplay between the pathways up-regulated in the Fast (‘Go’) and Slow (‘Grow’) phenotypes. Stress and DNA damage response, Src family activation and PI3K-Akt signaling in Fast cells displayed complex crosstalk with the elevated activity of ERK signaling and cell cycle regulation in Slow cells, with the casein kinase, protein kinase C and protein kinase A serving as important predicted signaling intermediaries. This modular organization allowed us to infer a simplified regulatory network common to both cell lines and reveal potential mechanisms of the network-level cross-inhibition between the parts of the networks corresponding to the Go and Grow phenotypes. Indeed, phenotype stabilization can be achieved through mutual suppression of phenotype-specific signaling and gene regulatory networks^30,31^, allowing for differential network activity patterns.

**Figure 5:**
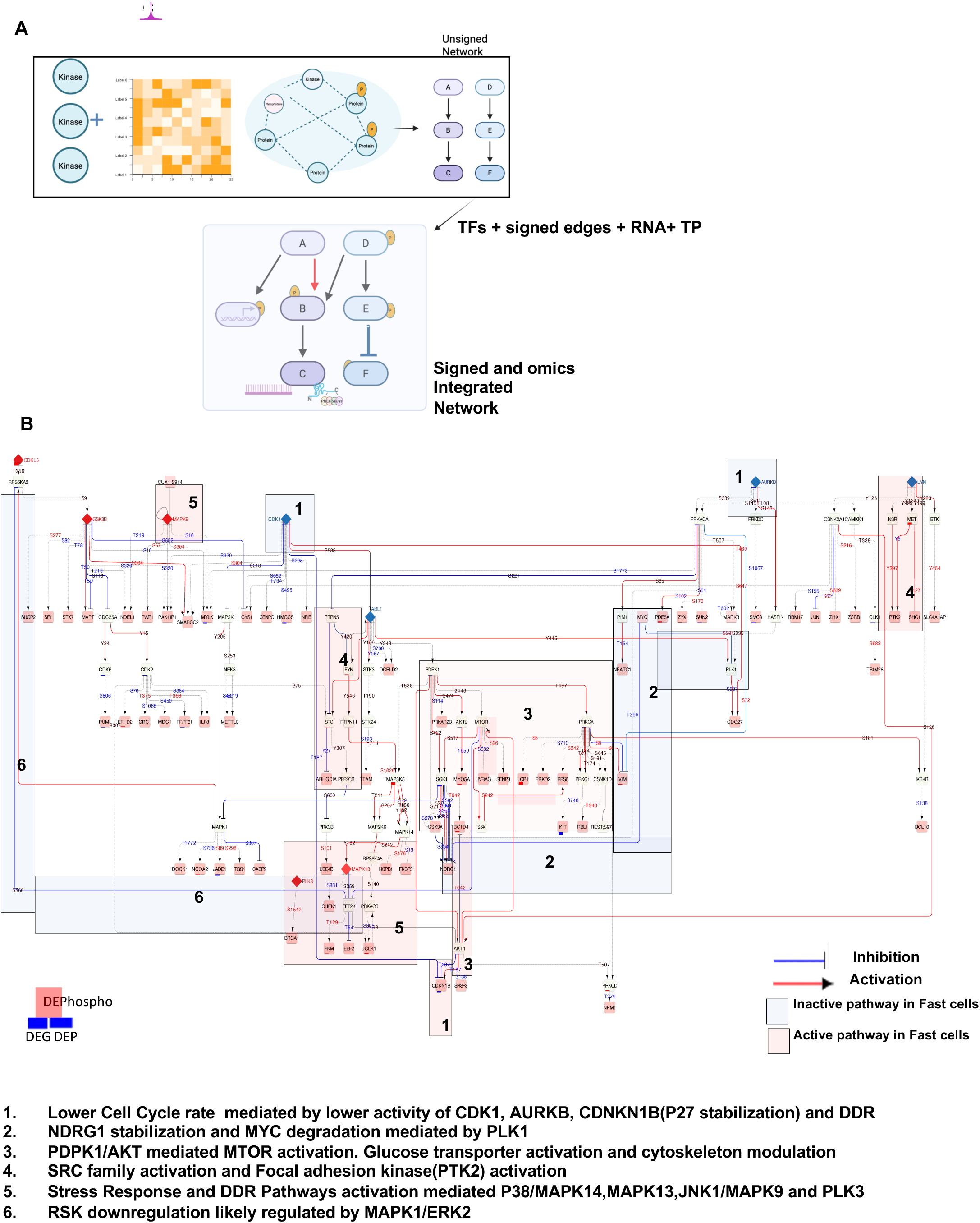
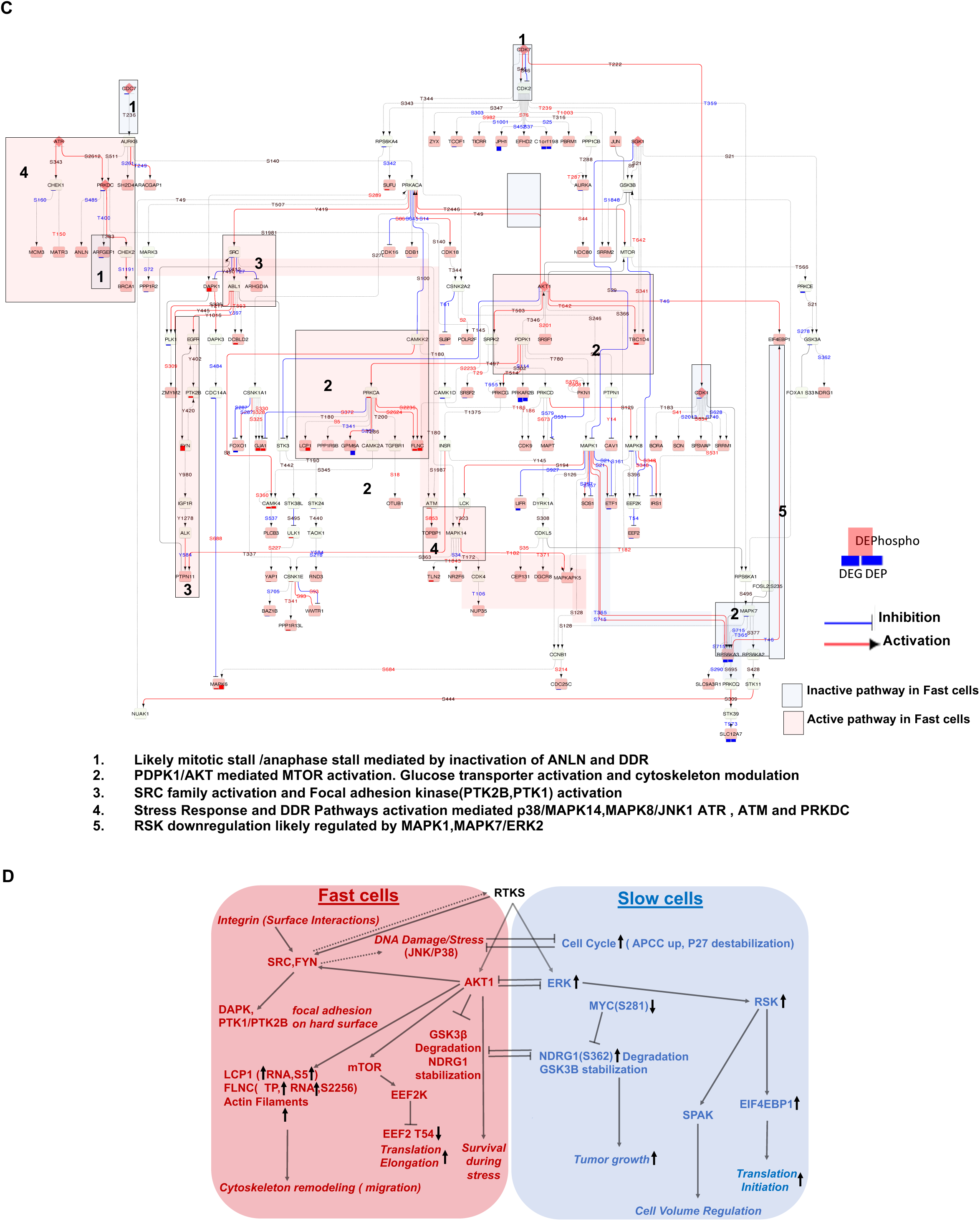
Distinct Signal transduction model for Fast and Slow Cells: (A) Network features from a directed unsigned network generated from PHONEMes were enhanced through the addition of Transcription factors, direction of regulation, missing edges, proteomics, and mRNA expression. (B and C): The directed signed signaling pathway that represents the difference between fast and slow cells as modelled by the PHONEMeS for A172 and T98G, respectively. Each pink node in the network is a differentially expressed phosphoprotein, and a green node is the filler phosphoprotein that connects the differentially expressed phosphoprotein. The diamond nodes are the significant kinases (Figure 3B and 3E). (D) Synopsis of figures B and C. Double negative feedbacks stabilize the fast and slow phenotypes.

One such interaction in A172 cells was suggested by the prior report that NDRG1 and GSK3B, which are both key signaling nodes in the inferred networks, can regulate each other through a double negative feedback, and that NDRG1 stabilization/GSK3B degradation was implicated in reduced GBM cell proliferation^32^. NDRG1 stabilization is regulated by MYC^33^, another inferred node in our analysis and the companion paper. Hyperphosphorylation of MYC at S281 mediated by PLK1 in Fast cells, observed in our study, suggests MYC degradation in fast cells leading to NDRG1 stabilization^34^. At the same time, fast cells in T98G show downregulation of MYC at the mRNA level, which can be a product of the interplay between the stress response and this key cell proliferation regulator, as further explored in the companion study. Another key double negative feedback we observed was the presence of double negative feedback between the DNA Damage/Stress response and the cell cycle. DNA damage activates cell cycle checkpoint responses and arrests the cell cycle in either G1/M or G2/M stage. At the same time, under normal conditions, the cell cycle kinases inhibit the checkpoint kinases that allow the cells to enter M-phase^35^.. Furthermore, we also observed the differential dominance of AKT and ERK in the Fast and Slow subpopulation. Fast subpopulation shows the dominance of AKT signaling, and the Slow Subpopulation shows the dominance of ERK signaling. The AKT1 and ERK signaling axis are parallel downstream targets of receptor kinases and integrin signaling. While these parallel signaling cascades overlap in terms of functional consequences, the dominance of one axis over the other depends on cellular context, localization, and the stimuli (cues)^36,37^..

Overall, this analysis paved the way for a detailed reconstruction of phenotype-specific regulatory networks based on high-throughput multi-omic data.

### Molecular Modeling Of Phosphosites Binding Domains Suggests Interaction With 14-3-3 as a Putative Functional Role Of Phosphorylation-Mediated Phenotype Specification

Phosphosites analysis may permit not only reconstruction of the signaling networks, but also assignment of functional significance by analyzing the likely domains and proteins binding to these phosphosites. We thus used PhosphoTune, an AlphaFold2-based model fine-tuned for protein phosphopeptide complex prediction^38^, to explore the resulting protein-protein interactions. Strikingly, we found that among the predicted binders to differentially phosphorylated proteins, a substantial proportion belonged to the 14-3-3 family (YWHAZ, YWHAQ, YWHAE,YWHAH,YWHAG,YWHAB, and SFN), which are known regulators of cell cycle, stress responses, and cytoskeletal dynamics.^39,40^ These predictions were validated with 14-3-3Pred, which specifically models 14-3-3-phosphoprotein interactions^41^.

We next explored whether the proteins containing phosphosites predicted to interact with 14-3-3 proteins belonged to functionally related classes. We noted that although most interacting proteins showed cell line specificity, there were essential overlaps both in the identity of specific proteins and their reported functions. For instance, in the Fast sub-populations of both lines, 14-3-3 was predicted to interact with TBC1D4 at T642 (Fig.6A and 6C), an interaction that is well established as essential for GLUT4 translocation to the plasma membrane^42,43^, supporting our network-based observation of AKT1 activation and downstream GLUT4 trafficking. Further relevance to phenotype-specific signaling was revealed by the identification of SPRED2 as a predicted 14-3-3 binding partner in Fast A172 cells (Fig. 6C). SPRED2 is a negative regulator of MAPK/ERK signaling, whose binding to 14-3-3 promotes ERK suppression and possibly RSK inhibition^44^. This is consistent with the ERK downregulation in Fast cells revealed via network reconstruction (Fig. 2B, 5B, 5C, and 5D).

**Figure 6:**
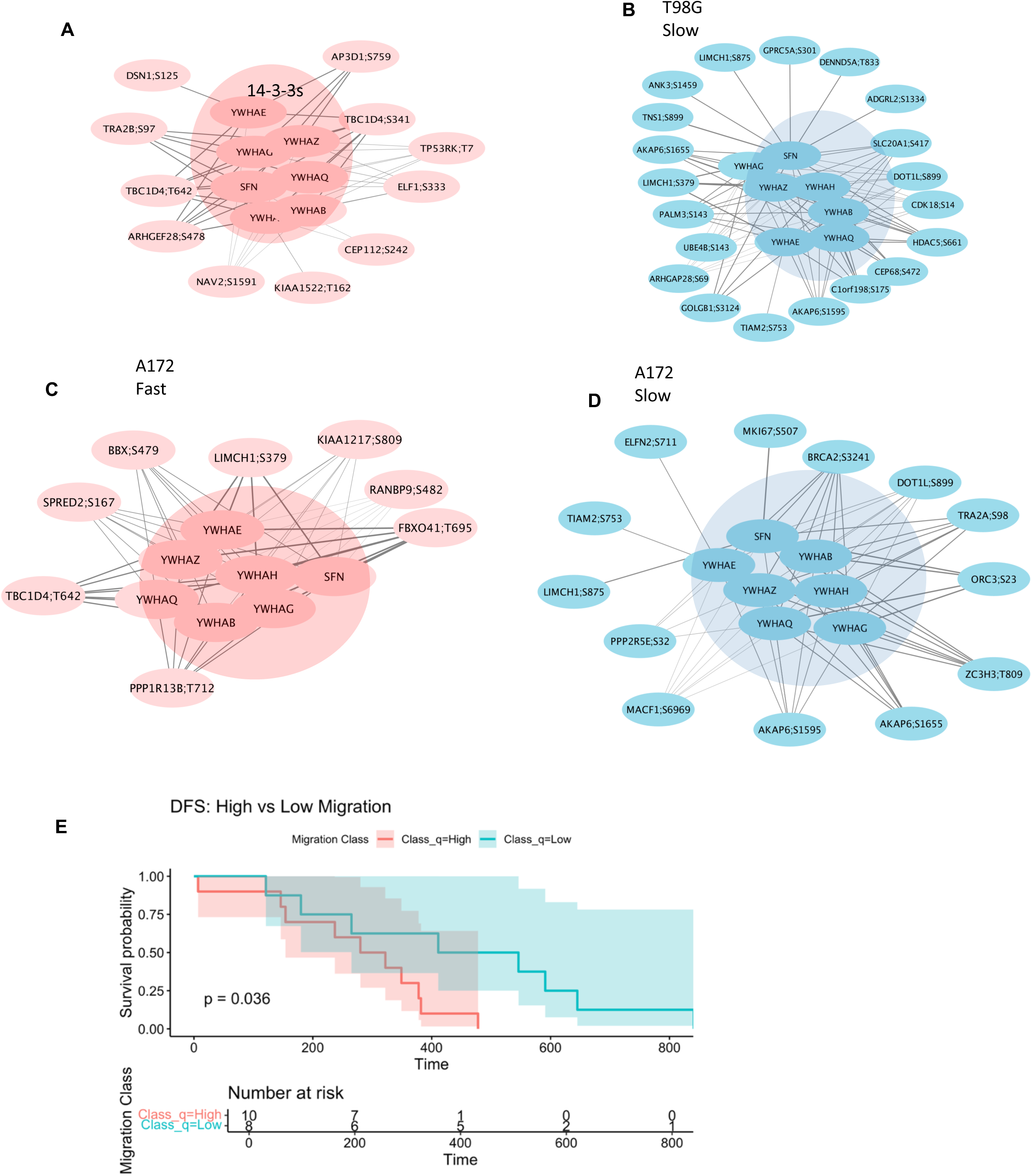
Protein motif-phosphoprotein interactions. (A, B, C, and D): Motifs (central highlight) predicted to interact with differentially expressed phosphoproteins in fast and slow cells. (E)Kaplan–Meier Disease Free Survival (DFS) analysis (quartile-based) for the patients stratified into high migration and low migration potential based on the network signature of the A172 cell line. Times are in Days. P-value is generated through the log-rank p-test.

Another groups of predicted 14-3-3 interactors were related to the control of stress response and cell cycle, consistent with the function of 14-3-3 and our prior findings of the importance of stress response for induction of cell cycle arrest and the Go phenotype. In particular, we found increased predicted 14-3-3 interaction in Fast cells of the following p53 regulatory proteins: PPP1R13B/ASPP1 in T98G and TP53RK in A172 (Fig. 6A and 6C), suggesting differential p53 pathway modulation between phenotypes through 14-3-3 binding. In particular, ASPP1 retains functional relevance in p53-mutant tumors, such as T98G, where it can still inhibit tumor growth^45^. ASPP1 function is localization-dependent: it regulates YAP/TAZ in the cytoplasm and modulates p53 transcriptional activity in the nucleus. Although 14-3-3 interactions with ASPP1 have not been reported, they might influence the localization-dependent roles. TP53RK phosphorylates p53 at Ser15 and enhances the transcriptional activity of TP53^46^. The interplay between cell cycle control and DNA repair regulation was suggested by predicted 14-3-3 interaction with PPP2R5E, BRCA2, and DOT1L in A172 and by DOT1L and CDK18 in T98G cells (Fig. 6B and 6D). CDK18 has been strongly implicated as a cell cycle regulator critical for appropriate response to replication stress ^47^. Apart from this role, CDK18 also negatively regulates the FAK/Rho signaling and actin dynamics^48^. 14-3-3 interactions may reinforce this function of CDK18, leading to the reduced invasion potential of the Slow subpopulation of T98G GBM cells. In A172 cells, a similar role may be played by BRCA2, which, in addition to its canonical role in DNA repair, can control the cell cycle. In particular, the phosphorylation of BRCA2 at S3291 detected here, mediating its predicted interaction with 14-3-3, signals mitotic entry^49^. PPP2R5E is a regulatory subunit of PP2A that directs its activity towards cell cycle regulators and MKi67, a classic proliferation marker, and origin recognition complex 3 (ORC3), which is found in actively dividing cells. Both these proteins are identified as the 14-3-3 interactors in Slow cells. Additionally, centrosome and spindle assembly control was suggested by CEP112 in A172 and CEP68 in T98G, with T98G also showing DSN1–14-3-3 interactions, potentially influencing kinetochore function and checkpoint signaling(Fig. 6B and 6D). Overall, these results indicate that differential 14-3-3-phosphoprotein interactions may contribute to the proliferative advantage of slow subpopulations by coordinating DNA replication, repair, and spindle assembly with cell cycle progression.

Of interest was the subset of interactions suggesting epigenetic control of gene expression. In particular, the analysis predicted HDAC6-14-3-3 interaction in Slow Cells T98G cells (Fig. 6B). On binding to 14-3-3, phosphorylated HDAC5 at S661^50^, it is held in an “inactive” state when it is present in the cytoplasm^51^. Therefore, this suggests that the deacetylase is inactive in Slow cells and active in Fast cells. NCOR1 interacts with both HDAC5 and 14-3-3 and stays inactive. Thus, the differential gene expression might be driven by the interaction of phosphorylated co-repressors with the 14-3-3 family.^52^ In addition, in both cell lines, 14-3-3 proteins were predicted to interact with DOT1L on S899 (Fig. 6B and 6D). DOT1L is an H3 histone methyltransferase involved in the regulation of diverse cell functions, including DNA damage response and the cell cycle.^53^ It is phosphorylated on S899 by a stress responsive p38β ^54^and, although this site has not been functionally annotated, its possible interaction with 14-3-3 might serve to control epigenetic regulation related to stress response in Fast vs Slow cells.

Finally, a large group of predicted 14-3-3 interactors were related to cell migration and cytoskeleton re-organization. In particular, two neuronal migration-related proteins were identified in Fast cells of both cell lines: NAV2 (S1591) in T98G and FBXO41 in A172 cells. NAV2 interaction with 14-3-3 has been implicated in neurite (axon) formation and possibly migration^55^, and even though the 14-3-3-FBXO41 interaction has not been reported in the literature, FBXO41 is also linked to neuronal-like migration^56^. These findings align with the concept that single GBM cells adopt neuronal-like migration programs, with 14-3-3 potentially acting as a regulatory hub^57^. This was further suggested by actin cytoskeleton remodeling hits that included LIMCH1, TIAM2 in A172, and LIMCH1, TIAM2, TNS1, and ARHGAP28 in T98G.

Together, these findings suggest that, functionally, a subset of phospho-sites differentially regulated within the pathways regulating cytoskeleton dynamics, cell cycle progression, DNA repair, and signaling pathways we identified within the Fast and Slow phenotypic states may work through interaction with 14-3-3 proteins. These interactions may control the spatial and temporal properties of the pathway regulations, in line with the prior work linking 14-3-3 proteins to response to various stressful stimuli and control of the cell cycle.

### Phosphoproteomics Signatures are Predictive of Disease-free Survival of GBM Patients

The phenotypes assessed in this study, i.e., the high proliferation and high migration, are both clinically relevant determinants of tumor growth and local invasion, followed by cell dissemination into the surrounding brain tissue. Therefore, it is of interest to examine if the phospho-proteomic analysis performed here might be potentially relevant for the prognosis of GBM progression in individual patients. Historically, the prognostic analyses have primarily been performed based on assessment of the transcriptomic and/or genomic signatures, as the phospho-proteomic data have not been as widely available. However, recently, such datasets have been collected in a patient-specific fashion. We thus assessed the clinical relevance of the phenotypic signaling networks identified in our analysis by exploring an independent global phosphoproteomics dataset collected from treatment-naïve glioblastoma patients, as deposited in Clinical Proteomic Tumor Analysis Consortium (CPTAC)^4^. To stratify patients based on migratory potential, we applied Gene Set Variation Analysis (GSVA)^58^ using two phosphopeptide sets derived from the A172 cell line network: one representing phospho-sites upregulated in Fast (invasive) cells, and the other in Slow (non-invasive, proliferative) cells (Fig.5B). Enrichment scores were calculated for each patient, and individuals in the first and fourth quartiles of the enrichment distribution were selected for survival analysis. Kaplan-Meier curves for disease-free survival (DFS) were generated by comparing patients who were tumor-free at their last follow-up (censored) with those who exhibited persistent disease or had died (events). We found that the patients classified as having phospho-proteomic signatures of low migratory potential showed a significantly higher probability of disease-free survival (Fig. 6E), highlighting the predictive power of the phospho-signature obtained through network reconstruction. This provided evidence that the signatures identified in this study from a limited number of cell lines can have more general applicability to GBM samples obtained in the clinical context.

## DISCUSSION

Current detailed characterization of phenotypic states of GBM cells is primarily based on transcriptomic and genomic profiling of the tumor samples. Although very useful, these efforts suffer from indirect inference of the post-translational mechanisms driving signaling networks and other key regulatory mechanisms underlying distinct cellular phenotypes. Here, we sought to address this persistent gap in knowledge by coupling phenotypic assays with the assessment of the multi-omics cellular profiles in two GBM cell lines, focusing on the phosphoproteomics analysis. The results strongly suggest that the Fast cellular phenotype associated with the high migration and slow proliferation (the ‘Go’ phenotypic state) is associated with increased PI3K-AKT-mTOR pathway activation and elevated DNA damage response, leading to promotion of cell migration, cell cycle arrest, and protection from stress-induced cell death. On the other hand, the Slow migration phenotype was associated with elevated proliferation (the ‘Grow’ phenotypic state), controlled by the ERK-RSK signaling pathway, stabilization of Myc, and related up-regulation of cell cycle-promoting networks. Based on this analysis, while also integrating the data from the proteomic and transcriptomic assessment of the matched samples, we could reconstruct the self-consistent signaling networks mediating distinct phosphorylation states corresponding to distinct phenotypes in both cell lines. This analysis further suggested that there might be mutually suppressive feedback interactions between the signaling sub-networks controlling each of the phenotypes, which might stabilize the phenotypic states. For instance, cross-regulation between NDRG-1, a stress-responsive gene up-regulated in Fast cells, and Myc in Slow cells can lead to the control of their stability and thus promote their mutually exclusive phenotype-specific expression and downstream regulation^32^.

A particularly striking result of our analysis is that the transcriptional factors regulating the transcriptomic programs distinguishing Fast and Slow phenotypic states are predominantly regulated post-translationally, through phosphorylation. This observation underscores the importance of assessing the phosphorylation more specifically and post-translational modifications more generally, to enable interpretation of commonly used transcriptomic data. This result also supports the critical role of phosphorylation-mediated signaling in modulating the gene regulatory networks and specifying distinct phenotypic states.

The persistent challenge in interpreting phospho-proteomic data is the lack of comprehensive annotation of the functional significance of the specific phosphorylation events. However, many of the functions endowed by this post-translational modification are mediated by the protein interactions involving phospho-site recognition domains. In our analysis, we used recently developed computational analysis platforms allowing prediction of such interactions based on the physically informed structural modeling. Our analysis strongly suggested that predicted interactors with the phospho-sites that displayed relative differential regulation in Fast vs. Slow cells predominantly belonged to the 14-3-3 family of signaling proteins. This finding was consistent with the known role of these proteins in mediating the stress response and regulating cell cycle, in various cell types, ^39,40^ which was also revealed by our analysis of phenotype-specific signaling networks controlling these processes. Further analysis supported these predicted interactions based on prior observations for a subset of implicated phospho-sites. This approach may serve as a useful intermediate in further analysis of the functional significance of large groups of currently unannotated phosphorylation outcomes.

Inherent in our methodology was the ability to assess the stability of the Fast vs Slow phenotypes in the presence of an ECM-mimicking cue that can serve as a strong inducer of mesenchymal phenotype across multiple cancers. Combining this cue with phenotypic filtering enabled by the RACE assay revealed that phenotypic stability led to a gradual enrichment of phenotypic and molecular level differences in T98G cells, whereas the A172 cells gradually lost their distinctive phenotypes and became a more uniform population dominated by the pro-migratory cue. Nevertheless, the distinct phenotypes, when detected, were controlled by similar networks in both cell lines and furthermore, the phospho-proteomic analysis allowed us to formulate a signature carrying prognostic significance for a cohort of GBM patients. More specifically, we were able to stratify the patients based on this signature into two groups that displayed significantly different disease-free survival times. This result suggested that the signaling networks associated with and possibly driving the Go-or-Grow switch in GBM cells may be universal across multiple cell lines and patient samples, and that the states of these networks can be predictive of clinical outcomes.

Overall, our results present an extensive characterization of the phosphorylation-dependent signaling and regulatory networks associated with two key aggressive phenotypic states in the most aggressive human brain cancer. This analysis not only can reveal the key underlying mechanisms driving these phenotypic states but may also serve as the basis for the development of new interventions targeting diverse populations of cancer cells. Indeed, many of the current and prospective drug development efforts are centered around human kinome and thus identification of the key signaling mechanisms driving clinically relevant cellular phenotypes can have immediate research and clinical relevance.

## Methods

### Rapid Assays for Cellular Enrichment (RACE)

The nanopatterned glass coverslip was laminin-coated (1 μg/cm²) at 37℃ for 2 hours. A stencil was placed perpendicular to the nanogroove direction at the midline of laminin coated nanopatterned coverslip. Cells were seeded into the stencil. For A172 cell lines, Accutase (Gibco) was used to dissociate the cells into single cells before seeding. Stencils were removed the next day, and fresh culture media was added. Cell culture media was changed every 48 hrs until 7 days. On day 7, the cells at both edges of the nanopatterned glass coverslip were collected as fast cells, and the cells in the midline of the device were collected as the slow cells. The sorted fast and slow cells were subjected to two more sorting round following the same protocol. After each rounds cells were expanded once (passage 1) for further omics data generations.

### Cell Migration Speed Measurement

Unsorted, Fast, and Slow cell subpopulations (10^4^ cells per condition) were plated on a laminin-coated nanopatterned device placed in 48-well plates, with three replicates per subpopulation. The patterned device was coated with 5*µ*g/ml of laminin at 37°C for 2 hours prior to cell seeding. Cell movement was recorded automatically using a 10X objective time-lapse microscopy for 12 hours at 10-minute intervals under controlled conditions (5% CO2, 37°C). Eight fields of view were recorded for each sample using Slidebook 6 software. The distance traveled (nm) by individual cells was tracked using the ImageJ tracking add-on MtrackJ^59^. We calculated the average migration speed, defined as the total distance traveled by the cells over the 12-hour period, the mean square displacement, and the trajectory plots^60^.

### Cell Proliferation Determination

Unsorted, Fast and Slow subpopulations are plated on two 96 well plates (three technical replicates for three biological replicates). We measured the ratio of luminosity of the cells from 24 hrs to 72 hrs using Cell Titer-Glo Luminescent Cell Viability Assay in the spectrometer at a wavelength of 560 nm. Cells were grown in 100 μL of medium to which the cell titer glo reagent was added at 24 h and at 48 h. The luminescence ratio at 72 hours/24 hours was measured as an indicator of cell proliferation. The p-values of the difference in proliferation were calculated, and the difference is deemed significant if the p-value <0.05.

### Mass spectrometry sample preparation and data processing

The cell samples were harvested, stored at -80 °C, and thawed before proteomic sample preparation. Cell lysis was performed using a 10 M urea buffer containing the cOmplete™ protease inhibitor cocktail (Roche, #11697498001). To ensure complete lysis, the tissue tubes underwent ultrasonic lysis by sonication at 4 °C for two cycles (1 min per cycle) using a VialTweeter device (Hielscher-Ultrasound Technology). The lysed samples were then centrifuged at 20,000 × g for 1 hour to remove insoluble material. Approximately 700 μg of protein was used for the subsequent digestion process. Reduction and alkylation were carried out using 10 mM dithiothreitol (DTT) for 1 hour at 56 °C, followed by 20 mM iodoacetamide (IAA) in darkness for 45 minutes at room temperature. The samples were then diluted with 100 mM NH₄HCO₃ and digested with trypsin (Promega) at a 1:20 (w/w) ratio overnight at 37 °C. The resulting digested peptides were purified using a C18 column (MaroSpin Columns, NEST Group Inc.). A 1 μg portion of the peptide sample was injected into a mass spectrometer for total proteome analysis, while the remaining peptides were used for phosphopeptide enrichment. The phosphopeptide enrichment process was performed using High-Select Fe-NTA kit (Thermo Fisher Scientific, A32992) following the manufacturer’s instructions^22^,^61^. Briefly, the resin of the spin column was aliquoted and incubated with 250 μg of total peptides for 30 minutes at room temperature, after which the mixture was transferred into a filter tip (TF-20-L-R-S, Axygen). The supernatant was removed by centrifugation. Subsequently, the resin-bound phosphopeptides underwent three washes with 200 μL of washing buffer (containing 80% acetonitrile and 0.1% trifluoroacetic acid), followed by two washes with 200 μL of water to remove nonspecifically adsorbed peptides. The phosphopeptides were then eluted from the resin twice with 100 μL of elution buffer (containing 50% acetonitrile and 5% NH₃•H₂O). All centrifugation steps were carried out at 500 × g for 30 seconds. The eluates were dried using a SpeedVac and stored at −80 °C before mass spectrometry (MS) analysis.

The samples were measured by data-independent acquisition (DIA) MS method as described previously^62–64^, on an Orbitrap Fusion Tribrid mass spectrometer (Thermo Scientific) coupled to a nanoelectrospray ion source (NanoFlex, Thermo Scientific) and an EASY-nLC 1200 system (Thermo Scientific, San Jose, CA). A 120-min gradient was used for the data acquisition at the flow rate at 300 nL/min with the column temperature controlled at 60 °C using a column oven (PRSO-V1, Sonation GmbH, Biberach, Germany). The DIA-MS method consisted of one MS1 scan and 33 MS2 scans of variable isolated windows with 1 m/z overlapping between windows. The MS1 scan range was 350 – 1650 m/z, and the MS1 resolution was 120,000 at m/z 200. The MS1 full scan AGC target value was set to be 2E6, and the maximum injection time was 100 ms. The MS2 resolution was set to 30,000 at m/z 200 with the MS2 scan range 200 – 1800 m/z and the normalized HCD collision energy was 28%. The MS2 AGC was set to be 1.5E6 and the maximum injection time was 50 ms. The default peptide charge state was set to 2. Both MS1 and MS2 spectra were recorded in profile mode. DIA-MS data analysis was performed using Spectronaut v17 ^65^ with directDIA algorithm by searching against the SwissProt downloaded mouse fasta file (date). The oxidation at methionine was set as variable modification, whereas carbamidomethylation at cysteine was set as fixed modification. For the phosphorylation data, phosphorylation (S/T/Y) (PTM score >0.75) were set as variable modification as well. Both peptide and protein FDR cutoffs (Qvalue) were controlled below 1% and the resulting quantitative data matrix were exported from Spectronaut. All the other settings in Spectronaut were kept as Default. In phosphorylation data analysis, the identification results were summarized with the PTM score > 0.75. At the same time, the quantitative table was filtered with PTM scores > 0.01 to minimize missing values due to phosphosite location for downstream analysis.

### Proteomics and Phosphoproteomics Normalization

Quantitative data matrix from MS for peptides/phosphopeptides was log2-transformed. Phosphopeptides expressed in at least two replicates per condition were retained. Among duplicated phosphopeptides, the one with the highest total intensity across all samples was kept. Missing values were imputed only if at least two replicates per condition contained values. If values were reported for one condition but missing in others, missing values were imputed using left-tail normal distribution imputation using PhosR package in R. The dataset was median centered after filtering. Batch correction was performed through normalization of samples using 100 stably expressed phosphosites across the samples^66^. The batch corrected samples showed reduced variability between biological replicates and increased difference between the samples (Figure S1-S2). Processed Data for Downstream Analysis: Only the filtered and normalized dataset was used in downstream analyses.

### Differential Expression Analysis

The differential expression between the samples were determined by fitting linear model through limma package ^67^. Phosphosites were considered differentially expressed if:

log2 fold-change *>* 1 or *<* −1

BH adjusted p-value *<* 0.05

### K means clustering of Conserved Phosphoprotein

Phosphoproteins (represented by unique phosphopeptides) that remained significantly differentially expressed across rounds were identified separately for each cell line (103 for T98G and 37 for A172). Prior to clustering, phosphopeptide intensity values were scaled, and Z-scores were calculated for each phosphopeptide across three rounds of RACE for Fast and Slow population. These Z-scores were then subjected to unsupervised K-means clustering in base R. The optimal number of clusters was determined manually based on inspection of hierarchical dendrogram branches and the overall clustering pattern. For each cluster, mean Z-scores were calculated to plot the long-term dynamics over rounds.

### Gene/Protein Centric Pathway enrichment

The log2 fold change rank-based Gene/Protein level Reactome Pathway^68^ Enrichment for total Protein, mRNA and Phosphoprotein were performed in R using PhosR^66^ Package. To perform the pathway enrichment of phosphoproteins at the protein level, multiple phosphosites mapping to the same protein were collapsed by selecting the phosphopeptide with the maximum absolute log2 fold change between the samples being compared.

### Site level-pathway enrichment and Kinase activity prediction

The activity of kinases was inferred from phosphorylation sites by performing Post Translational Modification (PTM) signature enrichment analysis (PTM-SEA)^19^. PTM-SEA uses the well-known single sample Gene Set enrichment analysis approach to infer the kinases from the database consisting of PTM signatures, ptm.sig.db.all.uniprot.human.v1.8.1.gmt. The peptides were ranked based on the log2(fold change), and following parameter was used to run the analysis. The database consists of Phosphosite-plus (PSP), drug based singatures, IKiP-db. We followed through the kinases that were predicted through PSP database only.

weight = 0.75,
correl.type = rank,
statistic=area.under.RES
nperm = 1000, min.overlap = 5,
global.fdr = FALSE

### Signaling Network Model for GBM subpopulation

Signaling networks for each GBM subpopulation were reconstructed using the PHONEMeS package in R^29^. PHONEMeS (PHOsphorylation NEtworks for Mass Spectrometry) is an integer linear programming (ILP)-based framework that infers signaling networks from phosphoproteomics data by integrating phosphorylation measurements with prior knowledge of kinase-substrate interactions. We implemented the upside-down model of PHONEMeS-ILP (PHONEMeS-UD), which enables bidirectional inference of kinase activity by considering both upstream and downstream signaling states. In PHONEMeS-UD, deregulated kinases inferred from phosphoproteomics data were integrated as putative perturbed nodes, allowing the model to infer both kinase activation cascades and phosphorylation-dependent regulatory mechanisms.

The **input** to PHONEMeS-ILP included:

1. Significantly different phosphoproteins between fast and slow populations from each round of the RACE assay.
2. Inferred significantly enriched Kinases from phosphosite-level data using PTM-SEA.
3. Prior Knowledge Network (PKN): Omni Path (filtered ProtMapper) to exclude low-confidence interactions.

### The following PHONEMeS-ILP parameters were used

Solutions=100
mipgap=0,
gap=0,
replace=2,
populate=5000,
intensity=4,
timelimit=3600,
penFac = 0.0001

**Outputs**: The output of PHONEMeS-ILP consisted of directed graphs mapping differential signaling interactions specific to each GBM subpopulation.

### Enhancements for Improved Network Reconstruction

To refine the inferred networks and enhance their interpretability, we implemented the following methodological improvements beyond the default PHONEMeS pipeline: **Addition of inhibitory and activating edges:** As the Phonemes is agnostic to the sign of regulation (inhibition and stimulation) we added the edges representing inhibition and stimulation when available using the PSP database. We incorporated regulatory interactions from PhosphoSitePlus (PSP) to distinguish between stimulatory and inhibitory phosphorylation events, ensuring a biologically accurate representation of kinase substrate interactions. **Logical consistency checks**: To prevent contradictions in inferred signaling pathways, logical consistency constraints were introduced. These checks ensured that inferred signaling did not contradict known biochemical regulation, such as activating phosphorylation leading to kinase inhibition. **Correction of edge connections:** We added any missing edges between the nodes in the network and master regulators of the input kinases in the network based on prior knowledge and literature.

### Late-stage omics Integration

To connect phosphoproteomics with gene regulatory mechanisms, we integrated transcription factor (TF) inferred from RNA-seq data into the phospho-signaling network. This integration was performed using both a **bottom-up approach** (linking TFs to upstream kinases) and a **top-down approach** (tracing kinase-driven regulation of TFs), resulting in a multi-layered regulatory network. The kinases regulating the TFs (bottom-up approach) were identified from the PSP-DB^20,69^ and the Kinase Library^70^. Additionally, we incorporated mRNA and total protein expression data to account for transcriptional and translational regulation of the differentially phosphorylated proteins. Specifically, we included genes with differential expression in total proteomics (adjusted p-value < 0.05 and |log₂FC| > 1) and mRNA expression (adjusted p-value < 0.05) to annotate the network further.

### Network Visualization

Cytoscape was used for network visualization^71^.

### Survival Analysis of GBM patients based on phosphoproteomics signatures

Phosphoproteomics data and corresponding clinical information were obtained from the Clinical Proteomic Tumor Analysis Consortium (CPTAC)^4^. To stratify patients based on migratory potential, we used Gene Set Variation Analysis (GSVA)^58^ to calculate enrichment scores for two phosphopeptides sets derived from the A172 cell line network (Figure 4B): one comprising upregulated sites in Fast cells and the other comprising upregulated sites in slow cells. The net migration enrichment score was defined as the difference between these two GSVA scores. GSVA was performed using the parameter mx.diff = FALSE, which computes enrichment based on the maximum deviation from zero, allowing for sensitive detection of partial or heterogeneous activation patterns within the gene set. Patients with positive net scores were classified as high migratory, and those with negative scores as low migratory. For survival analysis, patients in the 1st and 4th quartiles of the net enrichment distribution were selected. Kaplan– Meier curves for disease-free survival were generated by comparing patients who were tumor-free at their last recorded follow-up (censored) with those who had either persistence disease or had died by the last date of contact (events). Statistical significance was assessed using the log-rank test.

### Motif-Interaction Predictions

Phosphoproteomic data underwent initial filtering for significant changes in phosphorylation level (Log2FC > 2) and their statistical significance (P value < 0.05). Using annotated classes from the ELM database^72^, phosphosites were then to mapped putative short linear motifs containing filtered phosphorylated residues. To generate a list of potential phosphoprotein-protein interactions associated with differentiated phosphorylation, we used proteins containing domains from the PFAM/InterPro^73^ families corresponding to the mapped ELM classes. Kinases were excluded, as the PhosphoTune model used in the subsequent analysis is not applicable to them.

The potential phosphorylation-dependent protein interactions were then modeled using PhosphoTune - an AlphaFold2-based model ^38^specifically fine-tuned for protein-phosphopeptide complex prediction. The generated complexes were deemed confident if the mean pLDDT score was > 85 for the phosphopeptide and > 90 for the entire structure.

