## Supplementary figures and images for "ANALYSIS OF THE PHOSPHORYLATION NETWORKS CHARACTERIZING DISTINCT PHENOTYPIC STATES IN GLIOBLASTOMA CELL POPULATIONS"

### Supplementary Figure

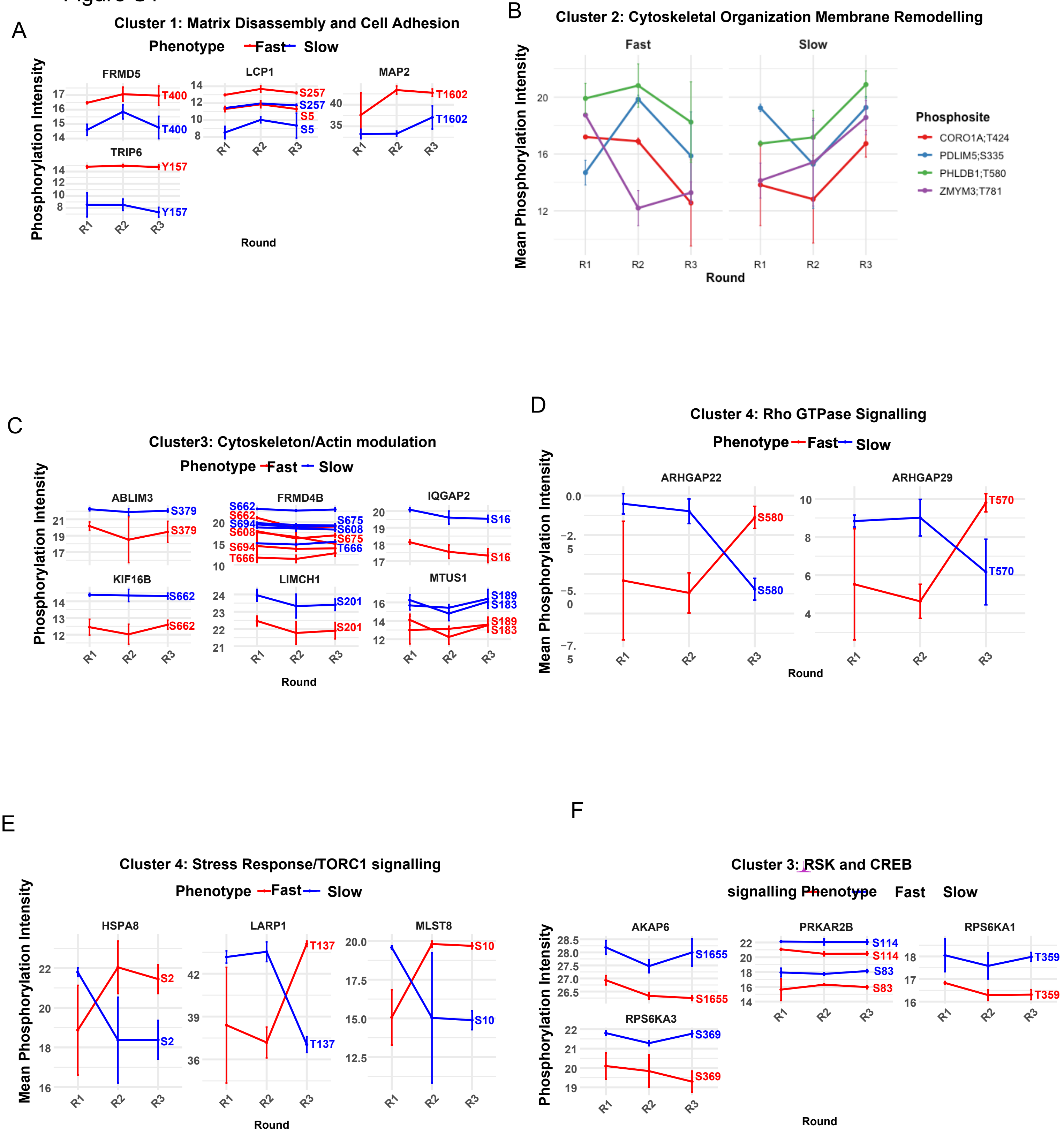

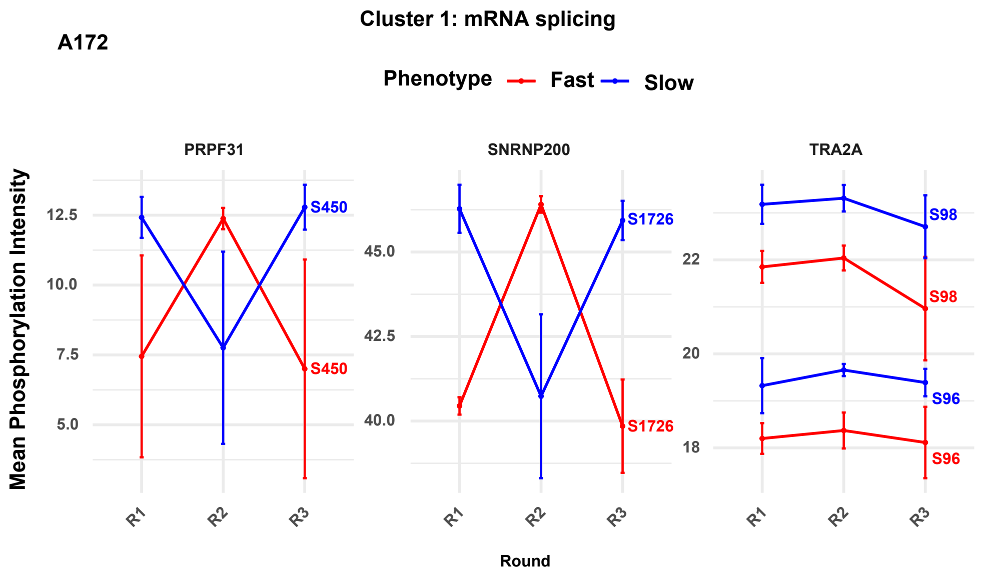

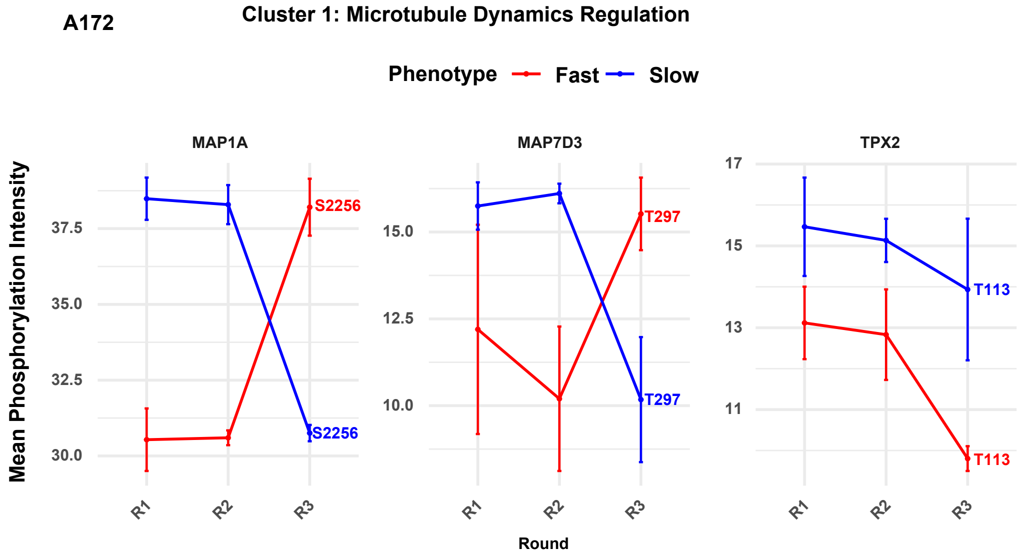

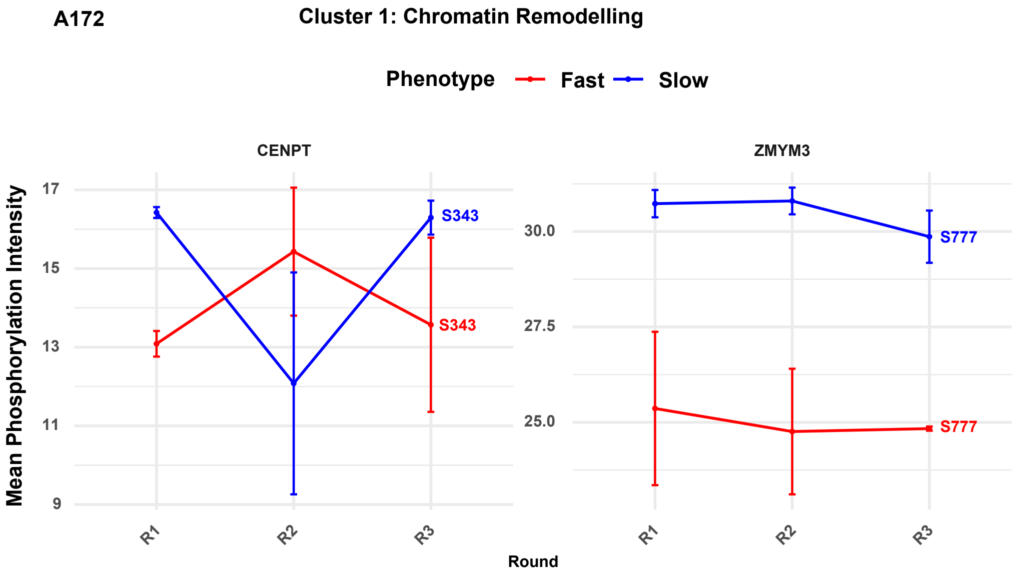

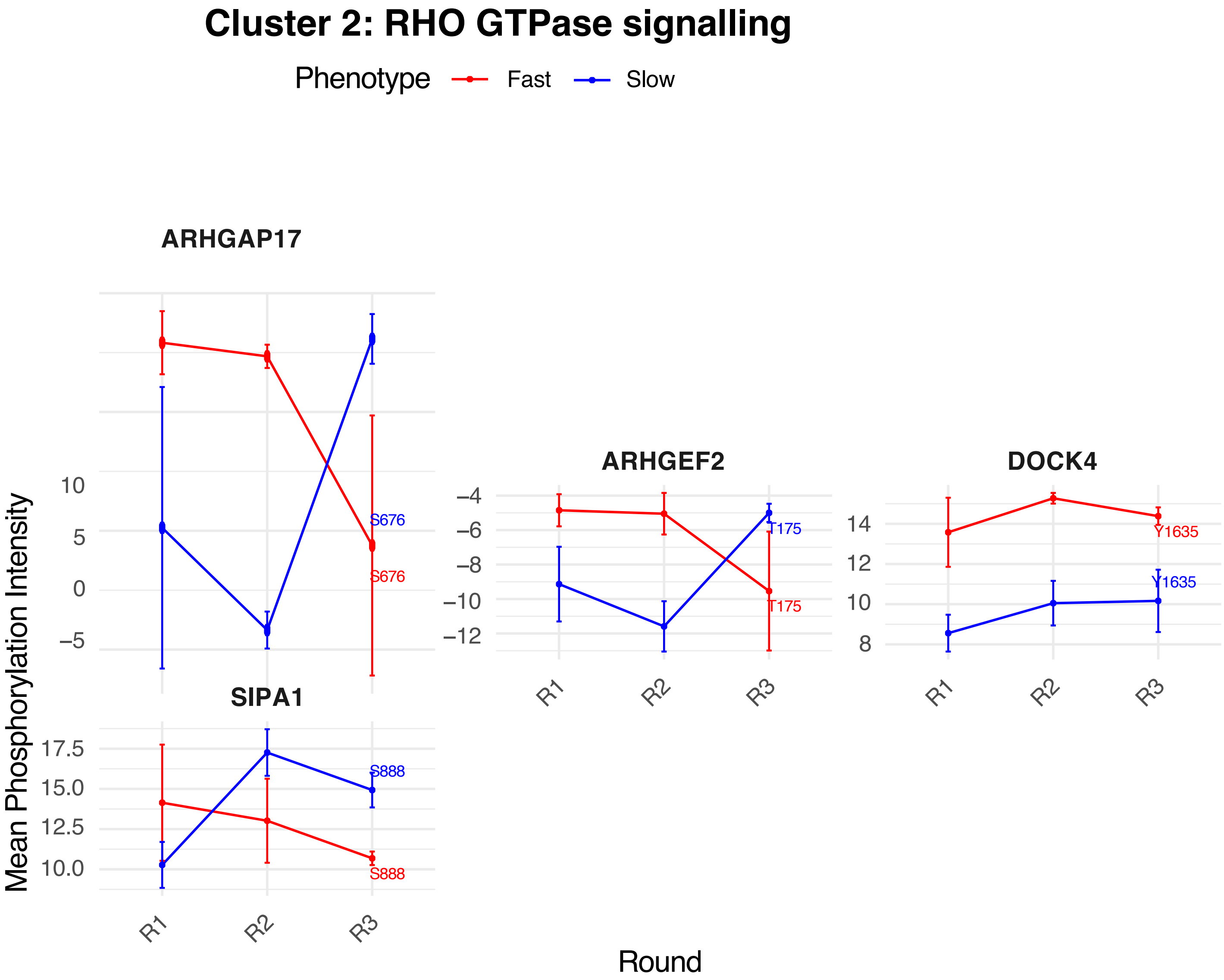
